# A SNP-based honey bee paternity assignment test for evaluating the effectiveness of mating stations and its application to the Ataun valley, Basque Country, Spain

**DOI:** 10.1101/2024.02.27.580467

**Authors:** Melanie Parejo, Egoitz Galartza, Jamal Momeni, June Gorrochategui-Ortega, Leila Farajzadeh, Jakob Wegener, Kaspar Bienefeld, Iratxe Zarraonaindia, Andone Estonba

## Abstract

Geographically isolated mating stations are deployed across Europe to facilitate controlled mating with selected drone-producing colonies. To assess the reliability of these stations, we developed a paternity assignment test using a custom Illumina genotyping chip with 6457 SNPs based on two metrics (number of mismatch alleles and kinship). The method demonstrated remarkable accuracy during validation with an independent dataset of known parent-offspring pairs, with an accuracy rate of 97.7%. We then applied the developed paternity assignment test to the *Apis mellifera iberiensis* mating station in the Ataun valley, Basque Country, Spain, in 2021. Drone-producing colonies in the valley were sampled and genotyped, as well as 156 worker offspring of queens mated at the station, and 56 drones collected in the drone congregation area. Out of the 156 worker samples, we could assign paternity of 120 (76.9%) to one of the drone-producing colonies in the valley, while 23.1% were of unknown patriline. Out of the 56 drones collected in the air, 52 (92.9%) were assigned to drone-producing colonies. We were also able to determine the colonies and apiaries that made the most significant contributions to the matings. This information aids in effective apiary management, including the selection of suitable mating station locations and the positioning of drone-producing colonies therein. Overall, our SNP-based paternity assignment test offers a valuable tool for evaluating mating station effectiveness across Europe, crucial for advancing breeding objectives in honey bee populations.

## 1. Introduction

Honey bees have a complex mating biology, with queens mating multiple times in flight away from their colonies, at so-called drone congregation areas (DCAs) (Koeniger et al. 2014). This makes their controlled breeding for conservation or genetic improvement programs quite challenging (Neumann et al. 1999a). For conservation purposes of local native stock, it is crucial to control mating in order to avoid hybridization with foreign honey bees (Parejo et al. 2016; Henriques et al. 2018a). For directed breeding programs, increased mating control and thus more accurate pedigree information ensures higher accuracy in estimating breeding values and ultimately leads to faster genetic progress (Plate et al. 2019).

To achieve controlled mating, various methods are employed. One approach involves setting up mating stations in geographically isolated locations, such as valleys or islands (Neumann et al. 1990b; Jensen et al. 2005). Geographic barriers help ensure that mating occurs with a certain degree of isolation, however, finding completely isolated locations is not straightforward given the high colony density in many regions (Chauzat et al. 2013) and the large mating flight distances of drones and queens (Ruttner and Ruttner, 1972; Jensen et al., 2005). Temporal isolation, another technique, involves controlling the timing of mating (Oxley et al. 2010; Musin et al. 2023). Artificial insemination is the most direct and controlled method, enabling total control over the genetic origin of the semen (Gillard & Oldroyd, 2020).

In Europe, numerous mating stations have been established to facilitate controlled honey bee mating (Bouga et al. 2011; Bienefeld et al. 2007). One such station is the Ataun mating station located in the Ataun valley, Basque Country, Spain, which was set up for the genetic improvement program of the native bee of the Iberian Peninsula, *Apis mellifera iberiensis*. The location was chosen because of its orographic characteristics and its scarcity of pre-existing apiaries. The mating station is managed by the bee breeding association ERBEL (www.erbel.eus) and has been running since 2019.

The effectiveness of the Ataun mating station and other stations alike must be evaluated to ensure that mating with the preferred drone-producing colonies is indeed controlled. Genetic monitoring tools that have been developed for estimating hybridization in *A. m. mellifera* (Parejo et al. 2016; Henriques et al. 2018a) and *A. m. iberiensis* (Henriques et al. 2018b) can give information on the percentage of pure matings. Observations of nuptial flights can also give information on the queen’s mating (Heidinger et al. 2014; Uzunov et al. 2023). However, a more direct way to evaluate the effectiveness of mating stations is the analysis of paternity between offspring of the virgin queens mated at the station and the drone-producing colonies. Paternity analyses aim to answer critical questions such as: drones from which drone-producing colonies have mated with the virgin queens, and what percentage of the mating drones are of foreign (uncontrolled) origin.

Traditionally, paternity in various species, including humans and animals, has been assigned using microsatellite markers (Pena & Chakraborty, 1994; Tian et al. 2008). While these markers are informative due to their multiallelic nature, they also have limitations: Their use is labor-intensive, tedious, and requires calibration among different laboratories (Ashley and Dow, 1994). In recent years, single nucleotide polymorphism (SNP) genotyping has become the standard method for assigning paternity (Clarke et al. 2014; Kaiser et al. 2017; Zhao et al. 2018). It allows for the simultaneous genotyping of thousands of loci, making it a more efficient and cost-effective approach. This shift to SNP genotyping is driven by the desire for greater accuracy, ease of use, and compatibility across different research settings. In honey bees, it has already been shown that SNPs outperform microsatellites in estimating hybridization (Muñoz et al. 2017; Parejo et al. 2018), and the use of SNPs to estimate kinship and pedigree information has already been performed in the context of genomic selection (Bernstein et al. 2022; Bernstein et al. 2023).

In this study, we present a paternity assignment test for honey bees based on SNP markers and built on the calculation of two metrics: (1) the KING kinship coefficient; and (2) the probability of identity-by-state equal 0 (P(IBS=0)). To this end, we employ the KING algorithm developed by Manichaikul et al. (2010) based on large-scale unlinked SNP sets that can be easily calculated between any pair of individuals. The KING kinship coefficient between a pair is defined as the number of minor alleles shared between two individuals, divided by the average number of minor alleles between the two individuals (Manichaikul et al. 2010). This metric for parentage testing has the advantage that it does not depend on population-estimated minor allele frequencies, *i*.*e*. the estimation of kinship coefficients is robust and independent of sample composition or population structure and thus reliable across different groups of individuals and populations (Manichaikul et al. 2010). This property, known as sample invariance, is important for ensuring that the method can be applied widely and accurately. The second metric, used in our approach, is the probability of identity by state equalling zero P(IBS=0). This metric measures the percentage of incompatible alleles shared between a pair of individuals, i.e. two individuals having homozygous opposite alleles. Unlike the kinship coefficient, this metric can differentiate parent-child relationships from sibling relationships.

Here, we developed a SNP-based paternity assignment test for honey bees based on the above-mentioned metrics and a honey bee genotyping chip with a total of 6457 SNPs. We first calculated kinship and P(IBS=0) on known parent-offspring relationships to set sensible thresholds for paternity assignment, then evaluated the performance of our method in an independent test set of known parent-offspring and unrelated pairs, and finally, applied the method to samples from the mating station in the Ataun Valley, Basque Country, Spain. The newly developed method proves valuable for optimizing apiary management, specifically aiding in the selection of appropriate mating station locations and strategic placement of drone-producing colonies therein. The SNP-based paternity assignment test holds great potential in assessing the effectiveness of mating stations throughout Europe, playing a pivotal role in progressing towards established breeding objectives.

## 2. Materials and methods

### Study area

The Ataun mating station is located in a narrow 10 km-long valley surrounded by mountains between 600 and 800m above sea level. The apiary of the nucs where the virgin queens are housed at is 330m above sea level, and it is situated at the opposite end of the valley entrance, at the most sheltered point (red star in Figure 1). Close by there are two apiaries with drone-producing colonies headed by sister queens from selected mothers of the breeding program of the ERBEL breeding association (www.erbel.eus): one with 6 colonies is located 150m from the apiary with virgin queens (apiary A, Figure 1), and another with 15 colonies is located at 2500m from the queens (apiary D, Figure 1). In addition, there are 3 other apiaries along the valley, one with two colonies at 650m (apiary B, Figure 1), one with 3 colonies at 2500m (apiary C, Figure 1), and one with 32 colonies at 7500m managed by a professional beekeeper member of ERBEL (apiary E, Figure 1). In the first two, the queens are changed every year with newly selected queens from the breeding program. The population in and surrounding the valley is composed of pure *A. m. iberiensis* bees.

**Figure 1.**
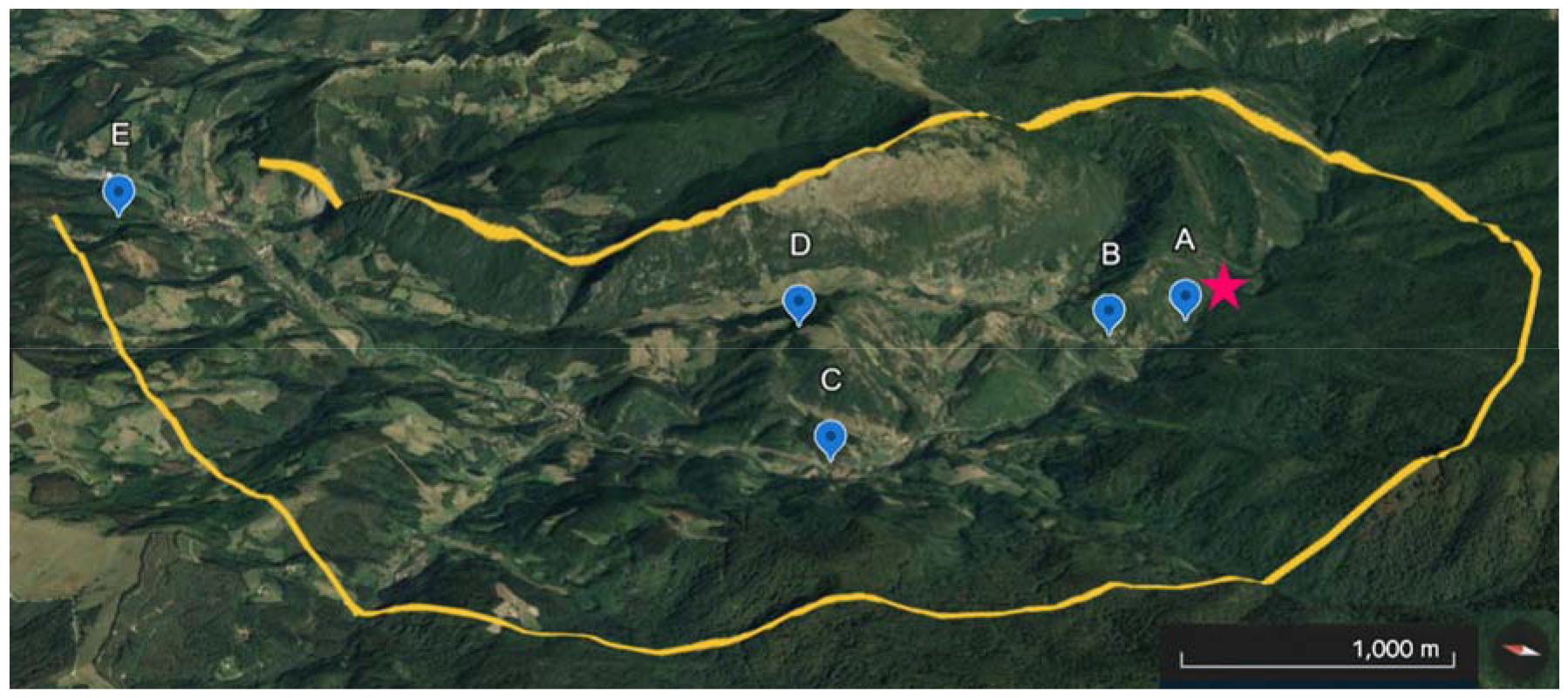
Map of the Ataun valley and its mating station. The red star indicates the location of the apiary where the mating nucs are placed. The blue placemarks with letters denote the apiaries with drone-producing colonies located along the valley. (Google Earth, accessed 03.12.2023).

### Samples of known relationships for threshold setting and validation

Iberian honey bee *(A. m. iberiensis)* samples of known parent-offspring relationship consisted of 6 queens (samples of 30 pooled drones to infer queen genotype) and 10-20 worker offspring each, totalling 99 (Table S1). Additionally, 6 Iberian queens (samples of 30 pooled drones to infer queen genotype) were sampled from an apiary outside the Ataun valley to be used for the inference of unrelated relationships to the mentioned 99 worker offspring.

### DNA extraction and SNP genotyping

DNA was extracted using sbeadex Livestock DNA Purification Kit (LGC, https://www.biosearchtech.com/). The samples were genotyped at a total of 6457 SNPs using the custom-made Illumina BeadChip array Infinium iSelect XT 96. The SNP chip includes 4165 SNPs for subspecies assignment (Momeni et al. 2021), as well as 2292 SNPs included in the same chip that have been designed for the selection of hygienic behaviour (SmartBees project, https://cordis.europa.eu/project/id/613960/reporting). The results were analyzed using Illumina’s GenomeStudio® software, and the genotypes of each sample were exported for further analysis.

### Kinship analysis

Genotypes were extracted from Illumina’s final report file using custom scripts and converted into plink .ped/.map files (https://www.cog-genomics.org/plink/1.9/formats). PLINK (https://www.cog-genomics.org/plink/1.9/) was then used to transform the .ped/.map files into binary plink files (.bed .bim .fam, --make-bed). Pair-wise relationship metrics for each putative parent-offspring pair (kinship and P(IBS=0)) were calculated with PLINK2 (--make-king-table, https://www.cog-genomics.org/plink/2.0/). The KING kinship coefficients are scaled such that duplicate samples have a kinship of 0.5, not 1, and first-degree relations (parent-offspring) correspond to a theoretical coefficient of ∼0.25. The probability of identity-by-state equal zero, P(IBS=0), reflects the percentage of mismatched alleles between a pair of individuals, *i*.*e*. two individuals having homozygous opposite alleles (AA vs GG). In an ideal case, this value is zero in a parent-offspring relationship and the appearance of mismatch alleles would exclude paternity. However, in reality, kinship and P(IBS=0) values deviate from the theoretical ideal case due to variable call- and genotype error rates among loci and individuals. Thus, the maximum proportion of genotyping errors must be determined empirically in a given population to set sensible thresholds for paternity assignment.

### Setting sensible assignment thresholds for A. m. iberiensis and evaluation of its performance

To set sensible assignment thresholds, we used 70% of the known-parent offspring sample pairs and known unrelated pairs (N= 304 pairs, training data), using queens and workers from the same colony, and outlier workers (see above). The distribution of the kinship coefficients and P(IBS=0) values of these known relationships were used to set the thresholds. Moreover, the distribution of P(IBS=0) values of drones and their colony of origin were considered for paternity assignments and threshold setting (Figure 2C). Due to the haploidy of drones (inexistence of minor alleles), it is not possible to calculate kinship following the formula of Manichaikul et al. (2010). Thus, for drones, we relied on the P(IBS=0) metric to assign paternity.

**Figure 2.**
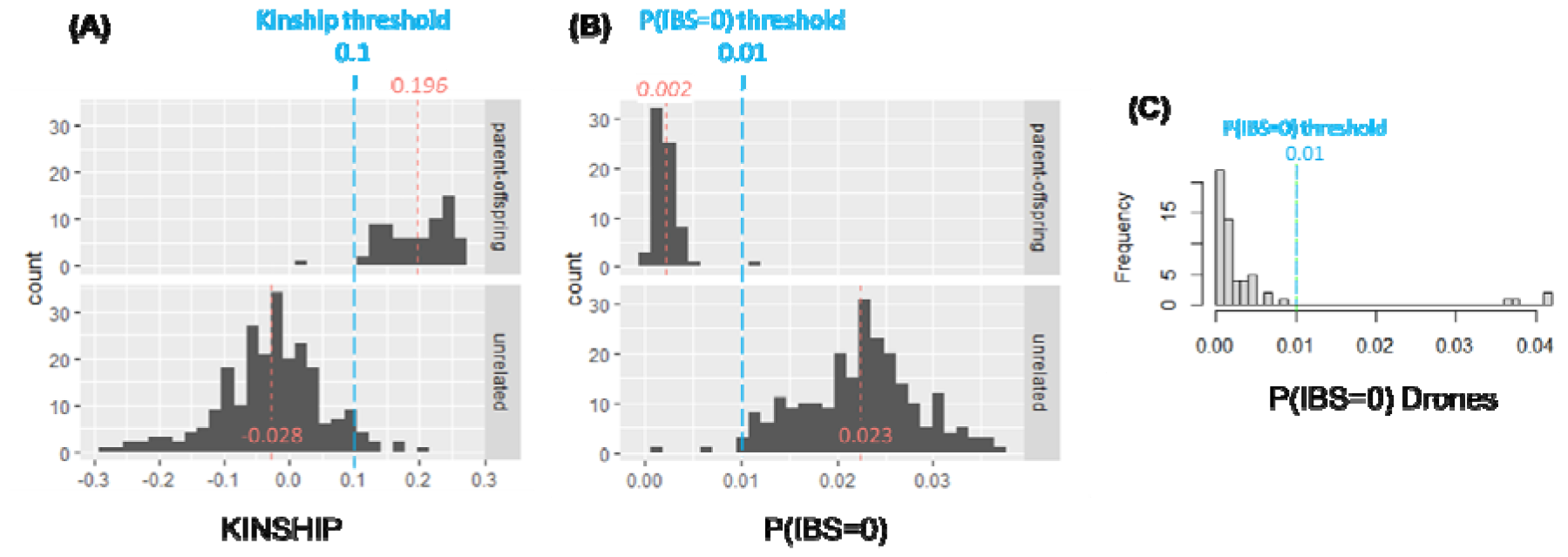
Threshold setting. Empirical distribution of the kinship coefficient (A) and P(IBS=0) value (B) of known parent-offspring and unrelated pairs (Train dataset, N= 304). The dashed red line indicates the group means, and the blue dashed line is the set assignment threshold. (C) Empirical distribution of the P(IBS=0) value of drones from the DCA and their most likely “father” colony. Considering a threshold of P(IBS=0), drones were assigned to their colony of origin if P(IBS=0)<0.01. Four drones with P(IBS=0)∼0.04 are considered drones with an unknown colony of origin.

Taking the 30% remaining dataset of the known relationships (N=131 pairs, test data), we evaluated the assignment performance of our method. To this end, we grouped the test data into the following categories: true positives (TP), *i*.*e*. pairs with a known parent-offspring relationship that were assigned parent-offspring according to our previously set assignment thresholds; true negatives (TN), *i*.*e*. unrelated pairs that were assigned an unrelated relationship; false positives (FP), *i*.*e*. unrelated pairs that were assigned parent-offspring; and false negatives (FN), *i*.*e*. parent-offspring pairs that were assigned as unrelated. Subsequently, we calculated precision (TP / (TP+FP)), accuracy ((TP + TN) / TOTAL), and recall (TP / (TP + FN)) in order to evaluate the assignment performance of our paternity assignment test.

### Application to the mating station of the Ataun Valley

We applied the paternity assignment test to the *A. m. iberiensis* mating station of the Ataun valley in 2021 in order to evaluate the effectiveness and reliability of the station. To this end, putative “father” colonies from the five apiaries along the valley were sampled (N=47; Figure S1). Eleven colonies (most of which from apiary E) that did not have drones at the time of sampling could not be sampled (Figure S1), but it cannot be excluded that they did produce drones at a later point. The sampled colonies (N=47) represent 81.0% of the total number of colonies located in the valley. Samples consisted of pools of 30 drone pupae each to infer the queen genotype of the colony. Moreover, in July 2021, a hot air balloon and queen mandibular pheromone were used to catch drones at the drone congregation area next to the mating station. In total, 56 drones were collected in the air, and they were also genotyped for paternity testing. Finally, 156 worker offspring were sampled in autumn 2021 and spring 2022 from 67 queens mated at the mating station (6 queens with 10 workers each, 4 queens with 9 workers each, and 61 queens with 1 worker each). Workers were sampled as pupae to make sure they were descendants of the mated queens and not drifters.

The DNAs of the samples from the Ataun mating station were extracted and genotyped as described above. With the raw genotype data, we then calculated kinship and P(IBS=0) metrics between all drone-producing colonies, *i*.*e*. the putative “father” colonies in the valley (N=47) and the 156 worker offspring of queens mated at the station. If the pairwise comparison of a given offspring with each of the putative “father” colonies did not pass the set thresholds, we deduced that none of the colonies fathered the tested worker. In the case that two or more “father” colonies pass the filters due to the relatedness of some of the drone-producing colonies, we assign paternity to the colony with the highest kinship.

We further tested whether the drones caught in the DCA next to the mating station originated from the drone-producing colonies nearby, whether they flew in from other apiaries along the valley, or whether they belonged to colonies of unknown origin. To this end, we calculated P(IBS=0) between all putative “father” colonies from the valley (N=47) and the 56 individual drones from the DCA using PLINK2 (--make-king-table). In the case, based on the set P(IBS=0) threshold value, that a drone is assigned to two or more “father” colonies, we consider the parentage given to the colony with the lowest P(IBS=0) value, *i*.*e*. the lowest number of incompatible alleles.

## 3. Results

### SNP genotyping

In total, for this study, 384 samples were genotyped at 6457 SNP positions. All samples were classified with high probability as *A. m. iberiensis* according to the subspecies diagnosis (Momeni et al. 2021). Samples included the pairs with known relationships, the drone-producing colonies of the Ataun valley, the worker offspring to be tested, as well as drones collected in the DCA. The mean genotype call rate per sample was 89.1% (Figure S2A). 113 SNPs had no call, i.e. genotyping call rate 0 (Figure S2B), indicating that the probes of these SNPs likely did not work in the *A. m. iberiensis* population. Another 1703 SNPs were monomorphic in our dataset, thus leaving 4641 informative SNPs for paternity analyses.

### Empirical distribution of relationship metrics of known relationships and set assignment thresholds

Kinship values of known parent-offspring relationships ranged from 0.013 to 0.262, with a mean of 0.196 (N=70), while for unrelated pairs kinship averaged -0.028 [-0.283 – 0.208] (N=234) (Figure 2A). Following this distribution, we set a boundary at 0.1 to distinguish parent-offspring from unrelated pairs.

The values of P(IBS=0) of known parent-offspring relationship ranged from 0.000 to 0.012 with a mean of 0.002 (N=70), and for unrelated pairs, P(IBS=0) averaged 0.022 [0.002 – 0.037] (N=234) (Figure 2B). Considering this distribution of P(IBS=0) of known relationships (Figure 2B), as well as the distribution of P(IBS=0) of the drones from the DCA with regards to the putative “father” colonies (Figure 2C), we set a threshold of P(IBS=0) <0.01 for the assignment of parent-offspring relationships in *A. m. iberiensis* bees.

Finally, for a higher reliability, we consider that both conditions (kinship>0.1, and P(IBS=0)<0.01) must be met for paternity assignment.

### Performance statistics

Given the established assignment thresholds, we evaluated the ability of our method to correctly assign paternity in samples of known ancestry using the independent test data set (N=131). The analysis revealed a precision of 93.3%, an accuracy rate of 97.7%, and a recall rate of 96.6%, indicating the consistent performance of our method in accurately identifying parent-offspring relationships from the test data set. The confusion matrix (Table 1) indicates that only one parent-offspring pair is misclassified as unrelated, and only two unrelated samples are misclassified as a parent-offspring relationship.

**Table 1.** Confusion matrix showing the number of correctly assigned and misclassified relationships.

|  |  | True relationship |  |
| --- | --- | --- | --- |
|  |  | parent-offspring | unrelated |
| Predicted relationship | parent-offspring | 28 | 2 |
|  | unrelated | 1 | 100 |

### Paternity assignment in the Ataun Valley

Application of the paternity analysis to the mating station of the Ataun valley in 2021 considering the set thresholds revealed that 120 out of the 156 worker offspring (=76.9%) could be assigned to their putative “father” colony, while 36 workers were considered offspring of an unknown colony. The mean kinship of assigned workers with their putative father was 0.216 with a range of [0.102 – 0.230]. The mean P(IBS=0) of assigned workers with their putative fathers was 0.003 ranging from 0.000 to 0.010.

The number of assigned workers per apiary can be seen in Figure 3A: Colonies from apiary A fathered 27 offspring which corresponds to 17.3% (=27/156) of tested workers and represents apiary A’s contribution to the effective matings. Apiary B had a contribution of 0.6%, apiary C 11.5%, apiary D 41%, and apiary E 6.4%. If we also consider the number of colonies per apiary, the average contribution of a colony per apiary is 2.9% for the six sampled colonies of apiary A, 0.6% for the colony in apiary B, 5.8% for the two colonies in apiary C, 3.4% for the 12 colonies in apiary D, and 0.2% for the 26 colonies in apiary E. When looking at individual colonies contribution, colonies B00000238 (apiary D), B00000200 (apiary C), B00000194 (apiary A) and B00000207 (apiary D) were the most reproductive with 15, 15, 14, and 14 assigned workers, representing 9.6% and 9.0% of the total matings, respectively (Table S2).

**Figure 3.**
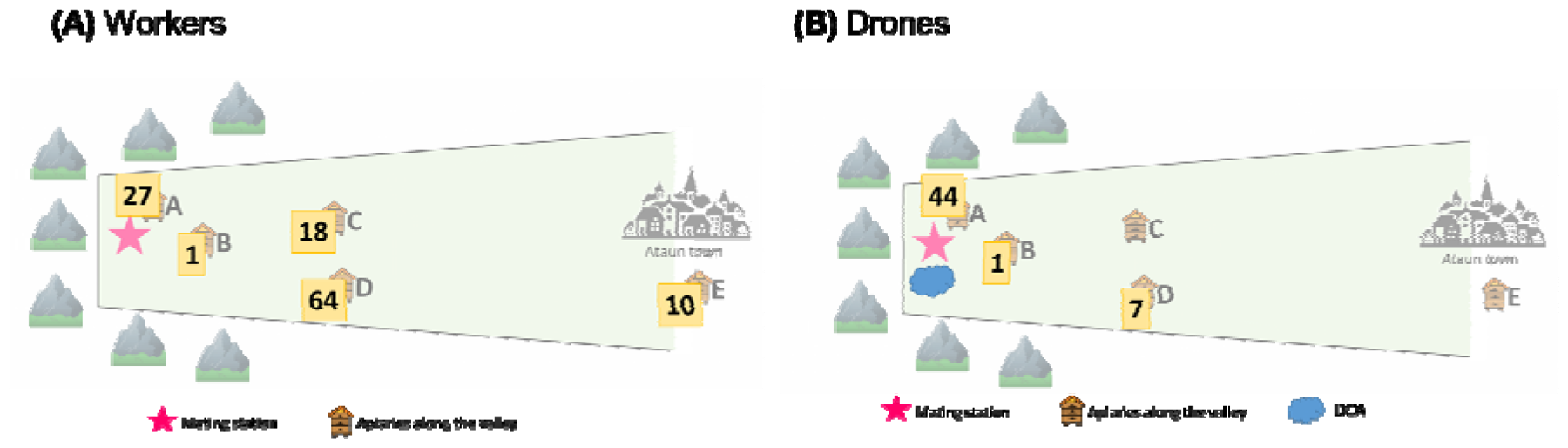
The number of assigned worker offspring (A) and drone offspring (B) per apiary. In the schematic figure of the Ataun valley the mating station is marked with a red star, the apiaries along the valley (A-E) are displayed by a hive, and the drone congregation area is represented by a blue cloud.

With regards to drones caught at the DCA next to the mating station, 52 out of the 56 tested ones could be assigned to their colony of origin (=92.9%). The mean P(IBS=0) of assigned drones with their colony of origin was 0.005 ranging from 0.000 to 0.009. Four drones were designated unassigned and had no known colony of origin, as indicated by their P(IBS=0)>0.01 reflective of an increased number of mismatch alleles (P(IBS=0)∼0.04 in Figure 2C). Forty-four (78.6%) of the drones originated from drone-producing colonies of apiary A, and only 1 and seven from apiaries B and D, respectively (Figure 3B; Table S3). Most of the assigned drones (20) belonged to colony B00000194 of apiary A.

## 4. Discussion

Mating stations are widely employed in Europe to guide virgin queens’ mating with selected drone-producing colonies. To assess the reliability of these stations, we developed a paternity assignment test using a custom Illumina genotyping chip with 6457 SNPs. The paternity of workers is determined based on mismatch alleles and kinship between worker offspring and potential “fathers” based on set thresholds inferred from empirical distributions of known relationships. Further, the P(IBS=0) values were also used to identify the colony of origin of drones. We show that our paternity test has a high accuracy and precision, and showcase its application to the *A. m. iberiensis* mating station of the Ataun valley in the Basque Country, Spain. This SNP-based test proves valuable for evaluating mating station effectiveness across Europe, crucial for achieving breeding goals.

The performance of our paternity assignment test was evaluated with samples of known ancestry (parent-offspring and unrelated relationships). The results unveiled an accuracy rate of 97.7% and a recall rate of 96.6%, emphasizing the method’s remarkable capacity to precisely identify parent-offspring relationships within the test data set. Only three relationship pairs were misqualified and the reasons thereof remain unclear. It is well possible that the two unrelated samples that were misclassified as a parent-offspring relationship, are more highly related (*i*.*e*. higher kinship value) simply by chance. However, it is to be noted that the achieved high performance of our method is found with *A. m. iberiensis*. The approach may have a different performance in other populations or subspecies with a different number of informative markers, and that will thus need to be evaluated empirically. Nevertheless, it is important to acknowledge that the KING kinship estimator utilized in our method offers a robust estimation of the relationship between any pair of individuals, regardless of the population structure (sample invariance) (Manichaikul et al. 2010). This implies that, given a sufficient number of informative markers, it is expected to perform equally effectively in substructured populations (e.g., European *A. m. mellifera* population) or potentially inbred populations (e.g., German *A. m. carnica* breeding population).

In a second step, we implemented our paternity assignment method to evaluate the effectiveness of the *A. m. iberiensis* mating station in the Ataun valley. In total, 76.9% of the worker offspring were assigned to their “father” colony, as well as 93.3% of the drones collected at the DCA were assigned to a drone-producing colony. Thirty-six workers and four drones were offspring of an unknown “father” colony, and several reasons could be thought of for these findings: (i) the presence of swarms or feral colonies in the valley, (ii) that foreign drones flew into the valley from neighboring valleys, or (iii) colonies present in the Ataun valley, but that were not sampled. In fact, it should be taken into account that some of the colonies located in the Ataun valley could not be sampled due to a lack of drones at the time of sampling (Figure S1). We assume that those colonies did not produce many drones, and thus, it is less likely they contributed to the matings. Nevertheless, it remains a possibility, and this could potentially explain the existence of the 36 worker offspring with unassigned paternity in our analysis. Therefore, it is reasonable to anticipate that the number of outlier drones in the Ataun valley is lower than the estimate presented in this study. Consequently, the effectiveness of the mating station is likely underestimated. Moreover, all samples were classified as *A. m. iberiensis* with a high probability, which highlights the mating station’s ability to guarantee pure-race breeding, even though we could not assign the totality of worker offsprings.

The insights derived from the paternity analysis not only offer an estimate of the overall effectiveness of the mating station but also enable us to pinpoint specific colonies and apiaries that made the most substantial contributions to the matings. Such information is very relevant for the strategic placement of selected drone-producing colonies. In the Ataun valley, the apiary that contributed most to the matings is apiary D (41%), followed by apiary A (17.3%) and C (11.5%) (Figure 3A). If we consider the average colony contribution per apiary, the order of highest contributions are colonies of apiaries C (5.8%), D (3.4%) followed by A (2.9%). These findings suggest that not only the colonies next to the virgin nucs must to be headed by selected queens from the breeding program (apiary A), but also the colonies at a distance, such as apiaries C and D located at 2500m from the mating apiary. This is in line with Tiesler et al. (2016) that in their book on honey bee selection recommend to set up selected drone-producing colonies at a certain distance for the organization and optimization of mating stations. Moreover, interestingly, in a study using microsatellites that analysed paternity in two mating valleys in England, it was found that most matings occurred with apiaries at the same distance (2500m) (Jensen et al. 2005). Ten workers (6.4%) were assigned to colonies from apiary E at the entrance of the Ataun valley, located approximately 8 km from the mating station. This is relatively low number considering that apiary E hosts a large number of colonies, more than the rest of the apiaries in the valley together, and supports the finding that the set up of the Ataun mating station is effective.

Furthermore, by the integration of data on the contribution of apiaries to matings with the insights gained from the paternity analyses of the drones captured in the DCA (Figure 3B), we can further speculate where in the valley the queens likely mated. Most of the drones caught in the DCA close by the mating station originate from apiary A (78.6%), however, apiary A contributed much less to the effective matings (17.3%). In contrast, seven drones of apiary D (12.5%) were in the DCA, but the apiary contributed much more to the effective matings (41%). From this, we can assume that a large part of the matings did not occur at the DCA next to the station, but that queens flew further downstream to mate with drones from apiaries C and D. The presence of at least another DCA in the valley seems thus likely. This is supported by direct observations of the duration of successful nuptial flights in the valley (personal communication from Egoitz Galartza). The finding that queens skip the nearest DCA is line with previous reports on mating behaviour (Koeniger et al. 2014).

It is to note that these findings, however, shed a light on the situation of the DCA and matings in the valley at that time. Other factors such as the number of drones per colony (Koeniger & Koeniger, 2007; Heidinger et al. 2014), geographical features, climatic and weather conditions (Galindo et al. 2012, Hayashi & Satoh, 2021) play a role in the formation and genetic structure of the DCAs and may be variable year by year or even in different times of the mating season (Jaffé et al. 2009, Koeniger et al. 2014). Nevertheless, results inferred from our developed paternity assignment, together with the direct observation of the nuptial flights of virgin queens (Uzunov et al. 2023), can deliver important information that contribute to optimizing the selection of suitable mating station locations and the positioning of drone-producing colonies therein. Ideally, to have a full picture of the effectiveness of the mating station it would advisable to conduct repeated analyses at different time points of the mating season offering insights into seasonal variations in honey bee mating dynamics.

The power of the test correlates with the number of polymorphic SNPs (Manichaikul et al. 2010), and the genetic diversity within the population. It is possible that the approach has a different performance in other populations or subspecies with a different number of informative markers, and that will thus need to be evaluated empirically. Moreover, likewise, the threshold established here for the *A. m. iberiensis* subspecies may not be universally applicable and should be re-evaluated for other subspecies. This limitation, however, could be overcome by increasing the number of SNPs, for instance by applying our approach to a more extensive SNP panel, such as the recently developed one for honey bees (Jones et al., 2022). In the original paper, Manichaikul et al. (2010) show via simulations that with 150k SNPs, the kinship coefficient estimation achieves optimal power to classify relative pairs up to the third degree.

Regarding the sampling design for assessing the effectiveness of the Ataun mating station, potential enhancements could be considered. For future studies, greater emphasis should be placed on sampling each colony in the valley to ensure comprehensive representation of all potential “fathers”. This approach would make clear that unassigned offsprings are from a mating with a drone from outside the valley. Additionally, the sampling strategy would involve selecting only one offspring per mated queen to ensure a reflection of distinct mating events, thereby avoiding the inclusion of repeated measurements (full siblings).

Another limitation in our approach and other methods of paternity assignment is it is unfeasible to determine the pure mating status of all queens in the breeding programme. Ideally, given the multiple mating of queens (Kerr et al. 1962; Adams et al., 1977), a large number of offspring per queen would allow us to determine the pure mating status of each queen and select only those for the next generation. Analysing this number of individuals per queen is economically unfeasible for routine purposes. Alternatively, a pool of worker bee offspring or queen spermathecal content (Yadró et al. 2023) could be tested, but pool data comes with genotyping and analytical challenges (Chen et al. 2022). To the best of our knowledge, it is not possible or technically and economically challenging to perform paternity analyses using pooled samples. Thus, while determining the pure mating status of each single queen is hardly possible, we estimated the percentage of matings with selected and foreign drones at the level of the mating station. Although this does not let us to predict which queen has mated exclusively with selected drones, we could use the gained information to optimize the safety and organization of Ataun mating station.

Finally, beyond its primary application in paternity assignment, the developed method holds promise for various additional uses. Through the genotyping of a substantial number of workers per colony, the test can be employed to estimate the number of matings and patrilines, shedding light on the mating frequency *i*.*e*. degree of polyandry within a colony or specific honey bee population. The analysis extends to exploring within-family relationships, elucidating the number of offspring sharing the same single drone father, originating from the same drone-producing colony, or having different fathers. Such comprehensive understanding of genetic relationships within colonies contributes significantly to our knowledge of honey bee reproductive dynamics. Moreover, this versatile tool also proves valuable in identifying and evaluating suitable locations for setting up new mating stations. By placing sentinel virgin queens in remote areas and subsequently evaluating the paternity of their worker offspring, it is possible to infer whether unknown drone-producing colonies contribute to the local DCA. These diverse applications underscore the method’s versatility and potential contributions to a comprehensive understanding of honey bee reproductive biology and population genetics.

