## Supplementary Material for "A SNP-based honey bee paternity assignment test for evaluating the effectiveness of mating stations and its application to the Ataun valley, Basque Country, Spain"

(2) ERBEL, Iberian Bee (Erle Beltz) Breeding Association, Zaldibi-Basque Country, Spain.

(3) Eurofins Genomics Europe Genotyping A/S, Aarhus, Denmark.

(4) Department of Biomedicine - Human Genetics, Aarhus University, Aarhus, Denmark.

(5) GrüBie Grüneberger Bienenzuchtgeräte Ashiralieva & Wegener GbR, 16775 Löwenberger Land, Germany

(6) Albrecht Daniel Thaer-Institute for Agricultural and Horticultural Sciences, Humboldt University of Berlin, 10099 Berlin, Germany

(7) Institute for Bee Research Hohen Neuendorf, Berlin, Germany.

(8) IKERBASQUE, Basque Foundation for Science, 48009 Bilbao, Bizkaia, Spain.

^†^ MP and EG have contributed equally.

**Supplementary Figures**

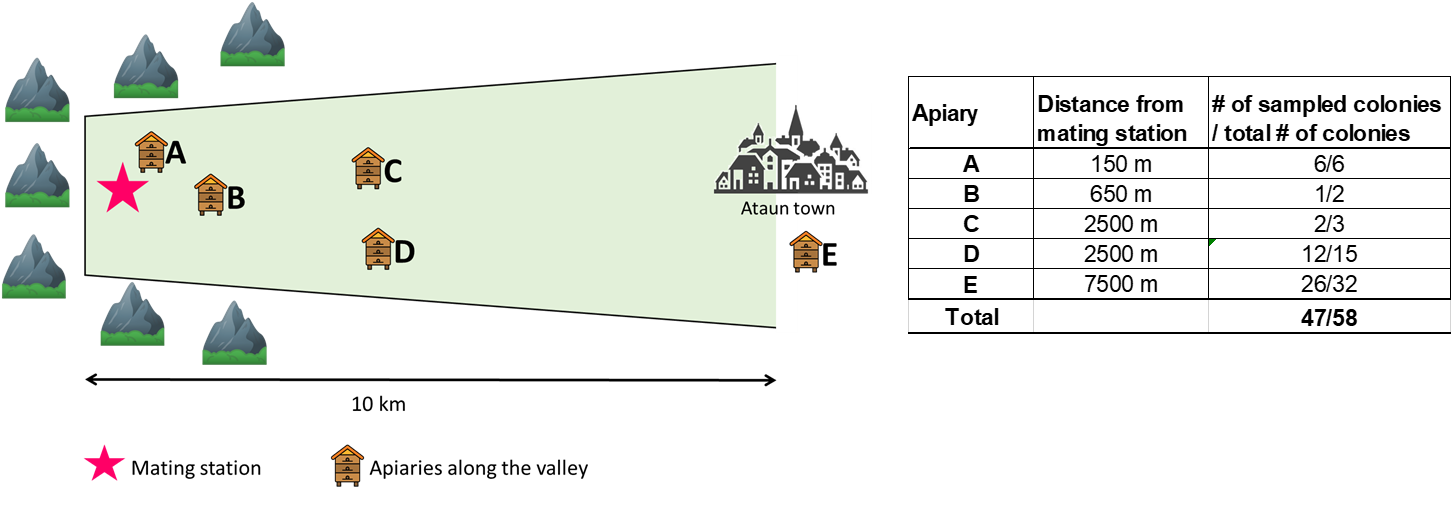

**Figure S1.** Schematic figure of the Ataun valley. The mating station is marked with a red star. The apiaries along the valley (A-E) are displayed by a hive. The distance of the apiaries from the mating station is shown in the table to the right, as well as the number of colonies sampled at each apiary and the total number of colonies at the apiary.

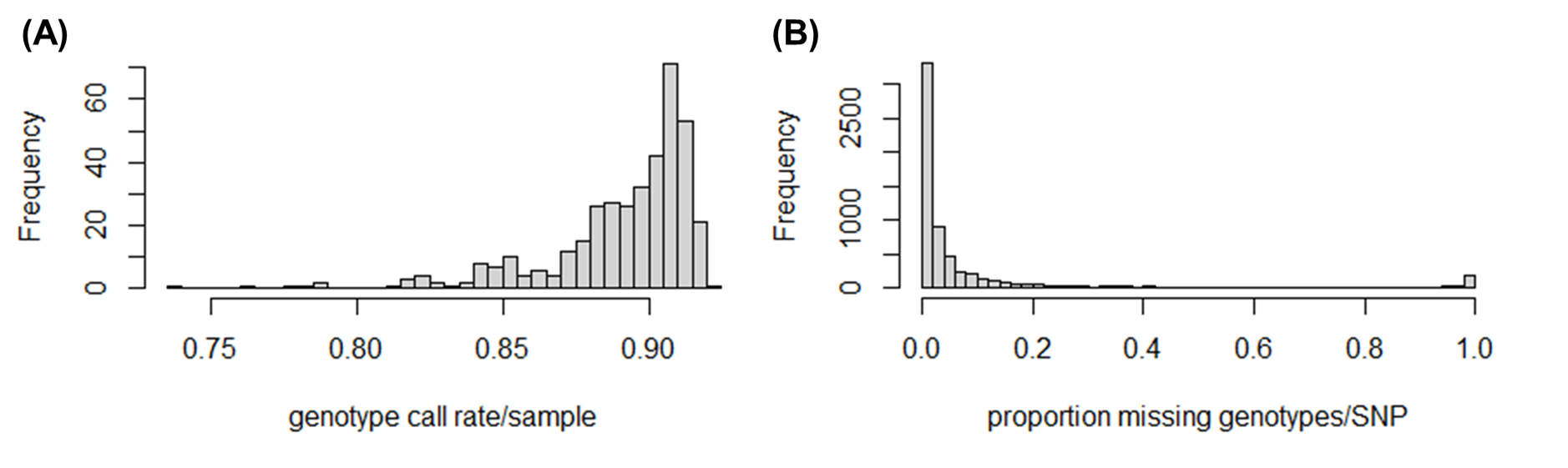

**Figure S2.** Distribution of genotype call rate per sample (A) and the proportion of missing genotypes per SNP (B).

**Supplementary Tables**

**Table S1.** Sampled queens and their worker offspring used to infer known parent-offspring relationships.

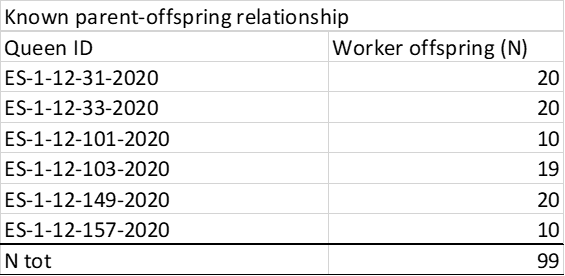

**Table S2.** The 20 drone-producing “father” colonies in the Ataun valley to which worker offspring have been assigned in 2021, totalling 120 worker offspring. The remaining colonies from along the valley were not assigned any paternity of the worker offspring analysed.

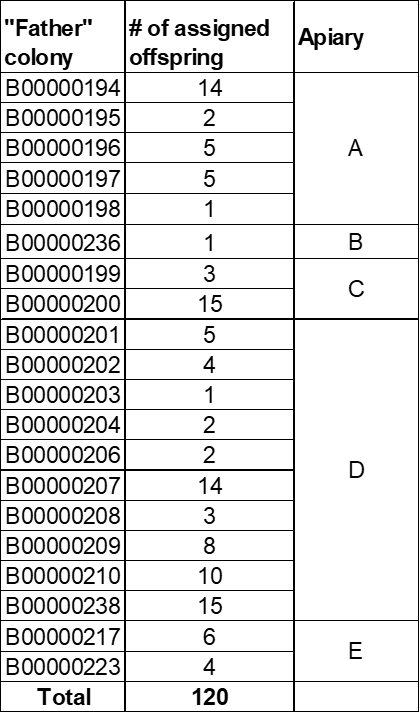

**Table S3.** The 11 drone-producing colonies of the Ataun valley that have been assigned drones caught in the DCA in 2021, totalling 52 out of 56 tested drones.

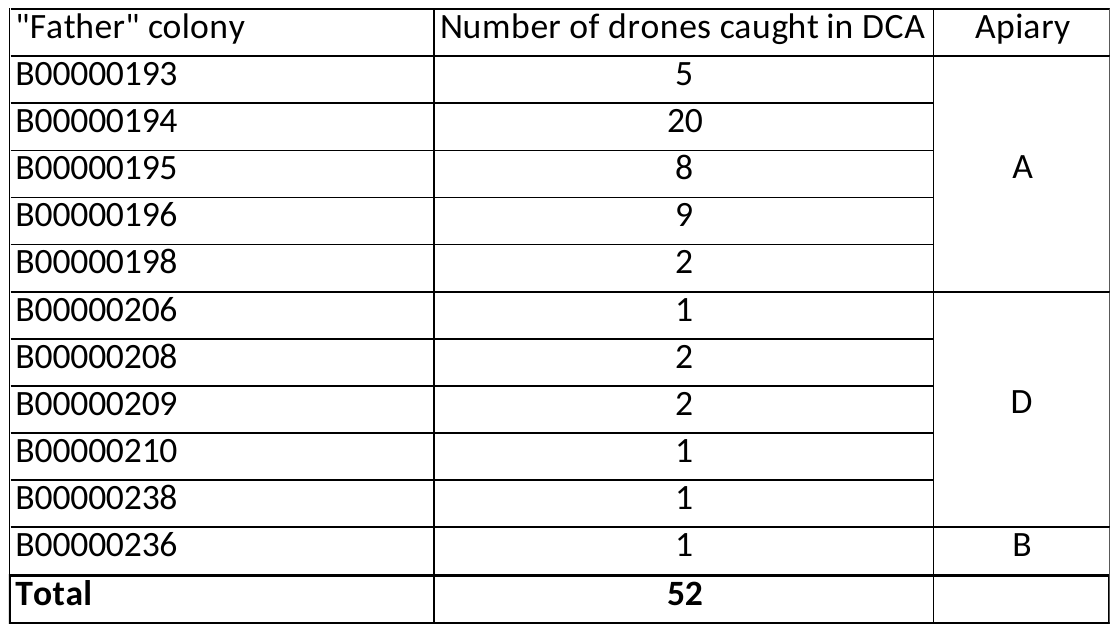
